# Constructing plasticity phenotypes to classify experience-dependent development of the visual cortex

**DOI:** 10.1101/2020.01.07.896191

**Authors:** Justin L. Balsor, David G. Jones, Kathryn M. Murphy

## Abstract

Many neural mechanisms regulate experience-dependent plasticity in the visual cortex (V1) and new techniques for quantifying large numbers of proteins or genes are transforming how plasticity is studied into the era of big data. With those large data sets comes the challenge of extracting biologically meaningful results about visual plasticity from data-driven analytical methods designed for high-dimensional data. In other areas of neuroscience, high-information content methodologies are revealing more subtle aspects of neural development and individual variations that give rise to a richer picture of brain disorders. We have developed an approach for studying V1 plasticity that takes advantage of the known functions of many synaptic proteins for regulating visual plasticity and using that to rebrand the results of high-dimensional analyses into a plasticity phenotype. Here we provide a primer for analyzing experience-dependent plasticity in V1 using example R code to identify high-dimensional changes in a group of proteins. We describe using PCA to classify high-dimensional plasticity features and use them to construct a plasticity phenotype. In the examples, we show how the plasticity phenotype can be visualized and used to identify neurobiological features in V1 that change during development or after different visual rearing conditions. We include an R package “v1hdexplorer” that aggregates the various coding packages and custom visualization scripts written in R Studio.

## 1. Introduction

Many neurobiological mechanisms regulate the development of the primary visual cortex (V1) and form a complex set of cellular and molecular states that can enhance or reduce experience-dependent plasticity. Often, studies of V1 development and plasticity focus on just a few of those mechanisms but the field is rapidly moving to large-scale studies. Studies are measuring tens to thousands of plasticity-related proteins or genes to understand how V1 develops (Benoit et al., 2015; Nowakowski et al., 2017; Siu and Murphy, 2018) and changes in response to disease (Smith et al., 2016; 2019) or visual experience (Beston et al., 2010; Dahlhaus et al., 2011; Majdan and Shatz, 2006; Tropea et al., 2006). Those big-data studies are beginning to reveal more subtle aspects of neural development and individual variations that give a richer picture of V1 plasticity. However, the complexity of those data is posing new challenges for understanding the combinations of mechanisms that underpin experience-dependent development and plasticity of vision. The idea of a *plasticity phenotype* is one way to tackle that complexity. The term plasticity phenotype has been used to describe the waxing and waning of gene expression during the critical period (Smith et al., 2019) but it has also be used as a tool to rebrand high dimensional protein or gene expression patterns into a reduced set of biologically meaningful features (Balsor et al., 2019). Here we present a primer for discovering suites of plasticity features, combining those features to construct a plasticity phenotype, and using the phenotype to classify normal and abnormal development of V1.

In this paper, we take advantage of insights gained from previous studies that identified the role of various neural proteins in V1 development and plasticity, especially glutamatergic and GABAergic receptor subunits, to select the set of proteins used for the examples in this primer (Cooper and Bear, 2012; Heimel et al., 2011; Hensch, 2005; Hensch and Quinlan, 2018; Maffei and Turrigiano, 2008; Smith et al., 2009; Yashiro and Philpot, 2008). Furthermore, because the approach uses protein expression, the same method can be applied to study V1 plasticity in both animal models and humans. The heuristic that we describe, a plasticity phenotype, will help for exploring and comparing neurobiological features that regulate experience-dependent development and plasticity of V1. The goal of constructing a plasticity phenotype is to transform the unique computational signature obtained from high-dimensional analyses of proteins and turn it into a biologically interpretable plasticity phenotype for V1.

We describe a workflow for constructing and using a plasticity phenotype by illustrating the steps in the statistical software R. Importantly, the steps include a visualization tool that enhances the exploration of the data. The data sets used in the examples are from studies by our laboratory of V1 development and changes after different types of visual experience on recovery after early monocular deprivation (Balsor et al., 2019; Beston et al., 2010). The examples address how to construct a plasticity phenotype, use it to discover clusters in the data, and apply the plasticity phenotype to identify biological mechanisms that underpin differences among ages or rearing conditions.

### Contributions of this paper

- We demonstrate how to combine measurements of plasticity-related proteins to construct and visualize a plasticity phenotype.
- We show how to combine the plasticity phenotype with high dimensional data analysis tools for partitioning data into clusters and identifying biological features.
- We illustrate how to use the plasticity phenotype to rebrand the data to discover biologically meaningful interpretations of the data.
- We aggregated the various packages and custom visualization code used in this paper into an R package “v1hdexplorer” that is available for download using the function install_github(“balsorjl/ v1hdexplorer”).

The rest of this paper is organized as follows. First, we review some of the high-dimensional data analysis methods that have been used in recent papers studying cortical development. Next, we present workflows with examples using PCA and tSNE and describe how to build and use a plasticity phenotype. Finally, we provide a brief summary and discussion.

### 1.1. Past work using high-dimensional analysis

#### Principal Component Analysis

The most commonly used high-dimensional analysis for exploring gene or protein expression in the brain has been principal component analysis (PCA) (Hotelling1933, 1933; Jolliffe and Cadima, 2016). PCA transforms the data, which is likely to include correlated genes or proteins, into a linear set of uncorrelated principal components that capture successively less of the variance in the data. Thus, individual cases can be visualized and analyzed in the transformed lower-dimensional space and that is often helpful for identifying clusters in the data. For example, a recent survey of human brain development used PCA to reduce the dimensionality of the data and identify differences among brain regions (Carlyle et al., 2017). Although this approach separated cerebellar samples from other clusters the unitless dimensions of PCA components made it hard to determine which biological features partitioned the samples into various clusters.

A different approach to using PCA takes advantage of known plasticity functions of a set of synaptic proteins and uses the basis vectors for each component (the weights for each protein) to attach biological significance to otherwise unitless dimensions (Jones et al., 2007). For example, the information from the basis vectors may reflect biological features such as sums of proteins, aspects of balances between pairs of proteins or maturational state of protein families that are known to affect plasticity (Beston et al., 2010).

#### t-Distributed Stochastic Neighbor Embedding

Another popular method for transforming and visualizing high-dimensional data is t-Distributed Stochastic Neighbor Embedding (t-SNE, (Maaten and Hinton, 2008)). tSNE measures the shortest distance between pairs of data points then calculates pairwise probability estimates of similarity across *all* dimensions. Often, these estimates are mapped onto 2-dimensional (2D) space by scaling the distance between data points and positioning similar data points closer together. The new mapping preserves local and global patterns thereby representing the relationships among data points to highlight clusters in the data. The artificial scaling makes it easier to identify clusters by either colour-coding points based on a known attribute (e.g. cortical area), or by applying a clustering method to the tSNE XY coordinates. Furthermore, the unsupervised nature of tSNE is particularly useful when exploring data without strong *a priori* knowledge of the biological features that may differ among the conditions.

A recent study of single-cell mRNA expression in the developing human brain used a combination of PCA and tSNE to analyze those data (Nowakowski et al., 2017). In that example, PCA was used to reduce the dimensionality of the data and tSNE to further reduce the dimensions and visualize clusters. This is a common workflow for analyzing and visualizing complex gene or protein data about brain development. Care is needed, however, when using the output from a PCA because the orthogonal components may not contain the information needed to partition the data into clusters (Chang, 1983).

Whether clustering is done with PCA, tSNE or some other method, the same challenge remains for studying brain development and plasticity -- how to link a holistic exploration of the data with the plasticity-related biological features that differentiate the clusters. The task of pinning down biological features is often done by sorting through the clusters using a large set of plots and univariate analyses aimed at finding individual proteins or genes that are over- or under-expressed in a cluster (Carlyle et al., 2017; Luo et al., 2017). That approach, however, loses sight of differences that arise from higher-order combinations of proteins or genes. To address that problem, we developed a new workflow for discovering combinations of proteins that represent high-dimensional features and a heuristic for analyzing the features that we call a plasticity phenotype. While the idea of brain phenotypes is not new (Cody et al., 2002), it has most often been used with brain imaging data. Our approach aims to construct a plasticity phenotype from neural protein expression data and use it to classify developmental and experience-dependent changes in V1.

## 2. Methods & Results

### 2.1. Note about the preparation of the data

Prior to beginning the analyses described in this paper, it is important to inspect and organize the raw data set. For example, if using Western blotting data ensure that the quantification of the bands did not include artifacts (e.g. bubbles, spots) or poorly labelled bands that could skew the results. Those data points should be omitted, and the missing data can be filled by imputation. A variety of imputation functions have been implemented in R and a package *impute* was developed for microarray data that imputes missing gene or protein expression data using a nearest-neighbour analysis (Hastie et al., 2019).

### 2.2. Description of the Example Data Set

The data set used for the examples in this paper comes from two animal studies of V1 development and plasticity (Balsor et al., 2019; Beston et al., 2010). The original data set has an *n*x*p* matrix comprised of *n*=567 rows of observations and *p*=7 columns of protein variables (Tables 1&2). The final matrix had 3,969 cells of data and after omitting 602 cells with poor labelling, the final number of data points was 3,367.

**Table 1.** Observations (*n*)

| Categories | Specific | Total |
| --- | --- | --- |
| Rearing Condition | Normal (9), MD (8), Reverse Occlusion (RO) (1), Binocular Deprivation (BD) (1), Binocular Vision (Long-term, LT; Short-term, ST) (5) | 24 |
| Regions | Central (2), Peripheral (8 or 10), Monocular (2) | 12 or 14 |
| WB Runs | 1,2 | 2 |
|  | **Sum** | **567** |

### 2.3. Constructing Plasticity Phenotypes to Describe V1 Development

The development of plasticity mechanisms in V1 is often described using scatterplots and curve fitting to capture the trajectory of protein or gene expression changes with age. For example, with the current data set that approach leads to 21 scatterplots and curves representing the 7 proteins and 3 sampling regions in V1. Even with the relatively small number of proteins in the example data set the number of possible trajectories grows quickly making it difficult to describe an overall pattern for the development of V1. Furthermore, that approach does not realize the potential of high-dimensional data since it is not inclusive of the full repertoire of proteins. Instead, holistic approaches that examine all proteins can identify patterns in the data that suggest how the biological functions might change. To address this combinatorial problem we developed a workflow that reduces the dimensionality of the data set (Fig. 1 A, B), explores and identifies biological features contributing to variance in the data (Fig. 1 C, D), validates the features (Fig. 1 E) and uses those features to construct a plasticity phenotype (Fig. 1 F).

**Figure 1.**
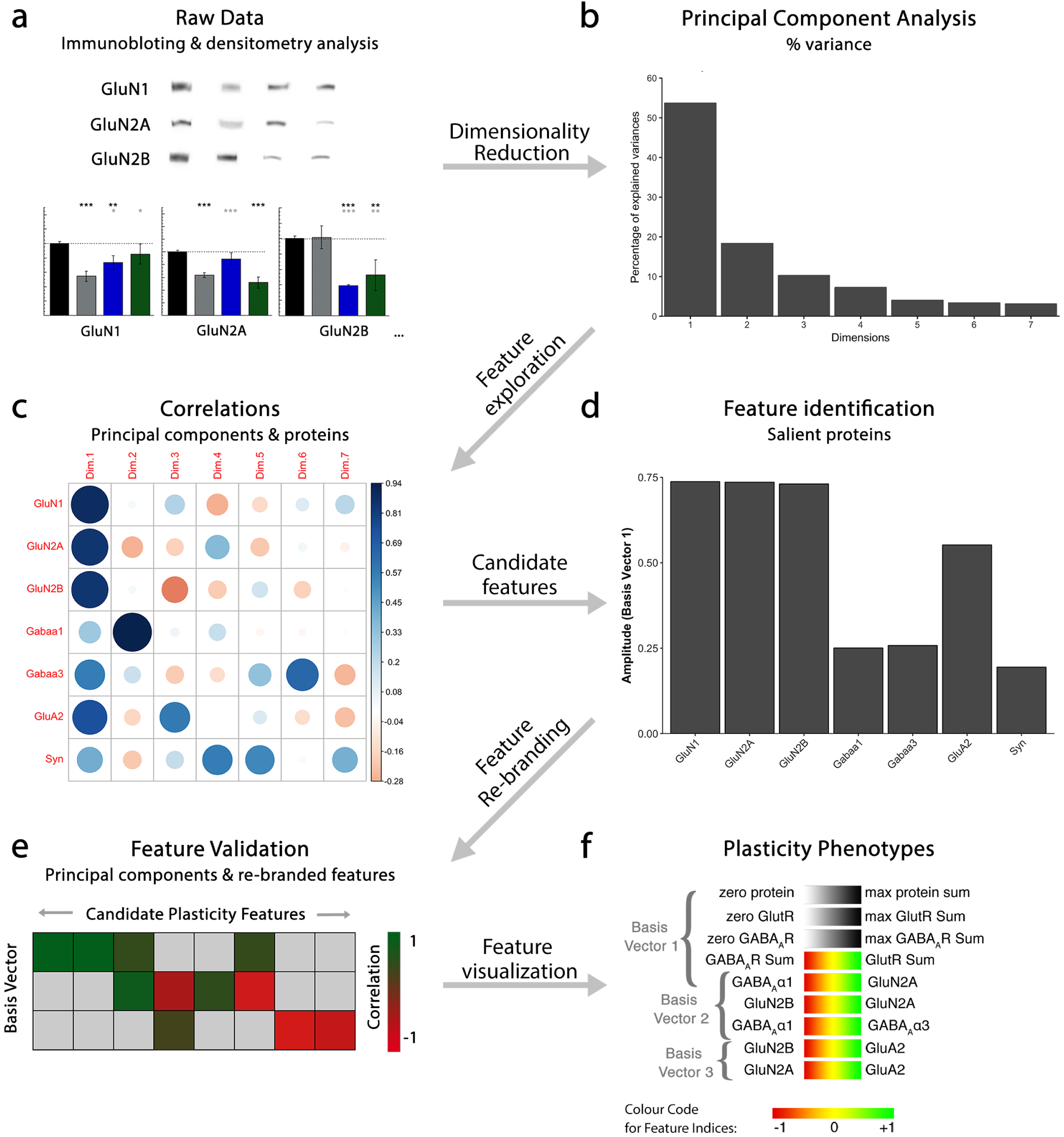
Analysis workflow to construct plasticity phenotypes. The analysis workflow for the data analysis described in the study. **a**. Immunoblots were quantified using densitometry, then comparisons among rearing conditions were made. Next, a series of steps were used to explore the data in a high dimensional space and create plasticity phenotypes. First, **b.** dimensionality reduction (principal component analysis) was done on the raw data, followed by **c.** feature exploration (correlations between principal components & proteins), **d.** identification of candidate features (saliency), **e.** feature re-branding (Correlation between principal components & features) and **f.** ending with visualization of those features by creating plasticity phenotypes.

#### 2.3.1. Dimension Reduction Using PCA

The first step in the workflow uses PCA to explore the high-dimensional nature of the data. We have implemented a two-step procedure that starts by reducing the dimensionality of the data and then uses the basis vectors for those dimensions to identify candidate biological features that capture the variance in the data.

A note of caution: many implementations of PCA do not work well when there are empty cells in the matrix. There are a variety of approaches that can be used, including imputation to fill in the empty cells, removing runs with missing data, or averaging across runs. In this example, we averaged protein expression across multiple runs of Western blotting.

This section is not an overview of PCA and we encourage readers to go to online tutorials to learn more about applying PCA to biological data. It is important, however, to emphasize that our use of PCA is a data-driven approach to understanding V1 development because *only* protein expression was used and no categorical information such as treatment condition, cortical area or age were included in the PCA.

This coding example (1) starts by extracting the columns (columns 10-16) from the dataset `my.data` that contained data for the proteins listed in Table 2.

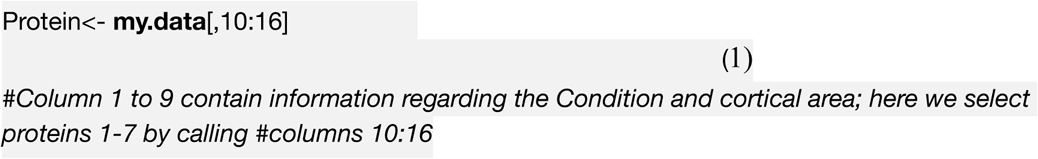

**Table 2.** Variables (*p*)

| Categories | Specific | Total |
| --- | --- | --- |
| Protein | Synapsin (Syn), GluN1, GluN2A, GluN2B, GluA2, GABA<sub>A</sub>α1, GABA<sub>A</sub>α3 | 7 |

The first step for performing PCA was to center and scale the data so that proteins with abundant expression did not obscure proteins with smaller but still significant variations in expression. Each protein was scaled and centered producing a standard deviation of ±1, and a mean of zero. Scaling data in R was done using the following example (2) of the base function *scale*:

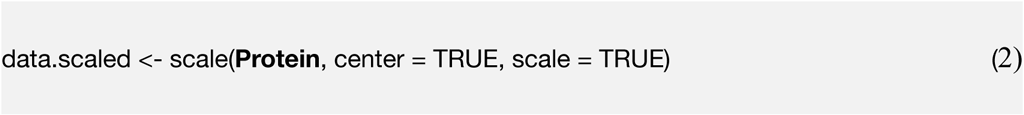

There are a variety of packages in R to do PCA, and here we used the *PCA* function in the *FactoMineR* package (Hudson et al., 2019; Lê et al., 2008). The *PCA* package produces eigenvalues and comes with excellent visualization tools to aid exploration of the relationships between principal components and features in the data set.

First, we ran PCA on the scaled data set `data.scaled` and saved the result as the object `pca`.

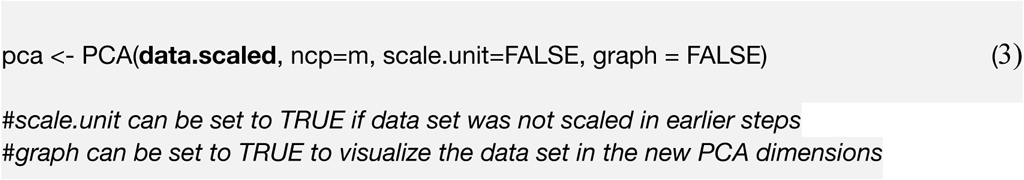

Principal components (PC) returned by this function are the set of orthogonal vectors in the object `pca` identifying the variance in `data.scaled`. The eigenvalues represent the magnitude of the variance captured by each PC vector, and the eigenvalue is largest for PC1 and successively less for each subsequent PC. An in-depth explanation of PCA and eigenvectors can be found here (Jolliffe and Cadima, 2016).

The eigenvalues for each PC were identified by consulting the `pca` object and stored as the `object` `eig.val`, as follows (4):

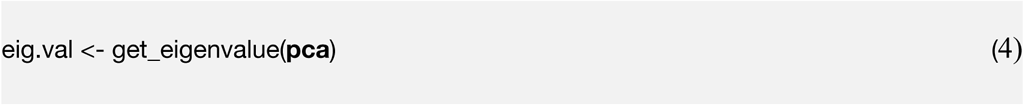

The first step of dimension reduction was to identify how much variance was captured by each PC, then rank the PCs from largest to smallest, and lastly, retain the set of PCs that capture a significant amount of the variance. Start with a Scree plot showing the amount of variance explained by each of the PC dimensions. The following coding example (5) demonstrates how to consult the pca object to create a scree plot.

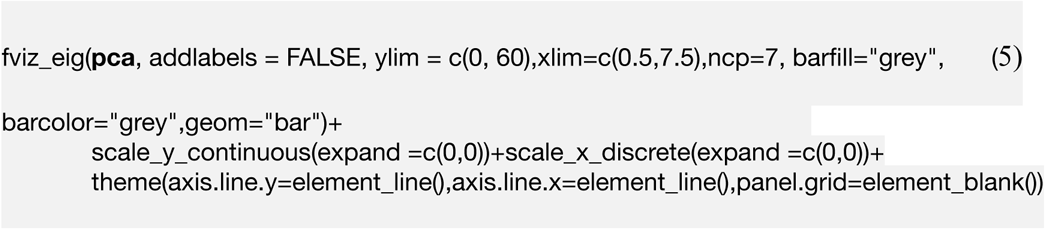

The Scree plot (Fig. 2A) shows the decreasing magnitude of the variance explained by the 7 PC vectors. A variety of methods have been used to identify significant dimensions (Hoyle, 2008) and here we used the simple rule that retains successive dimensions until the amount of cumulative variance explained was ≧ 80%. In this example, Dim1-3 explained 82% of the variance and those were used in the next steps to identify candidate plasticity features.

**Figure 2.**
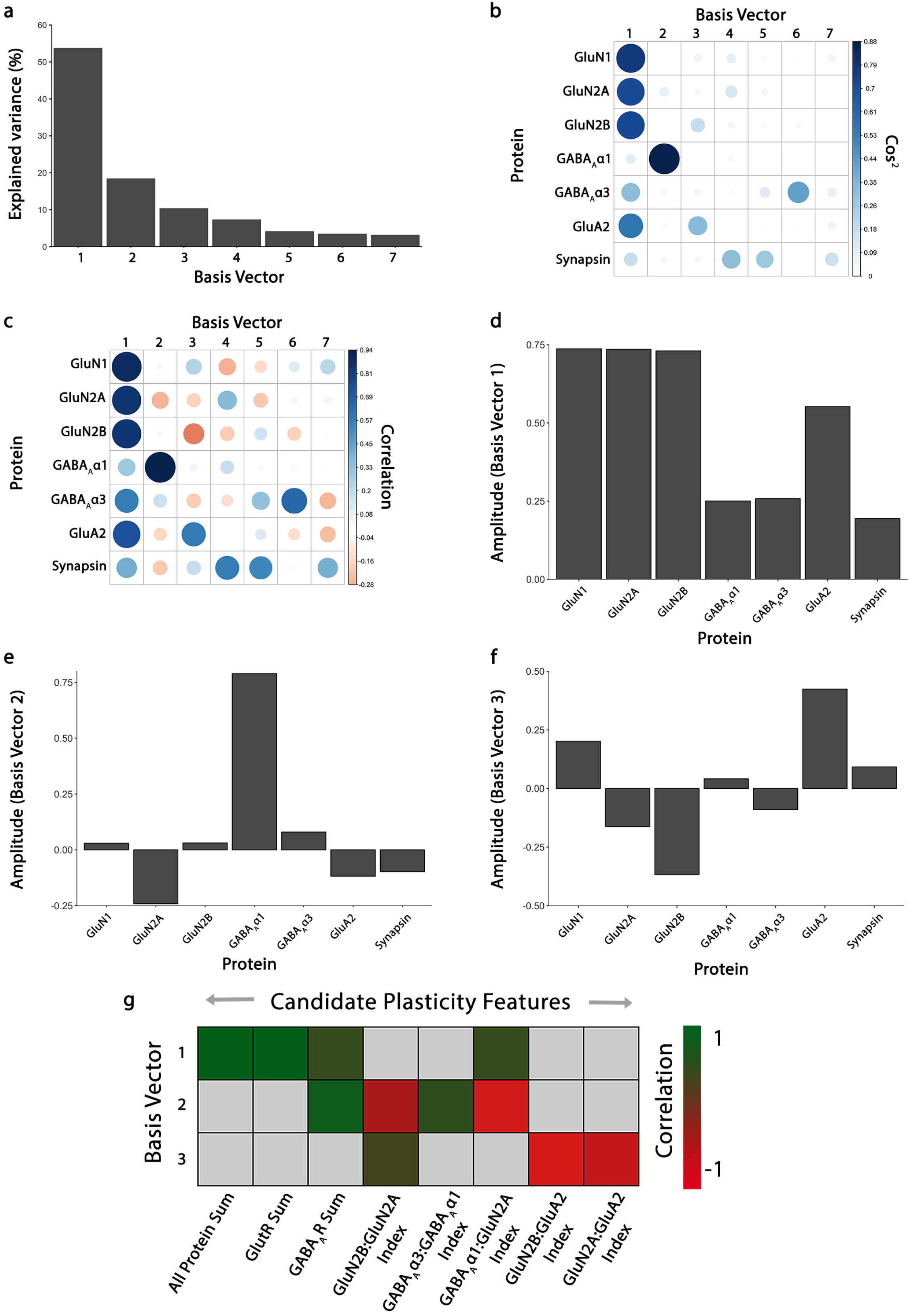
Using principal component analysis to identify candidate plasticity features. **a**. The explained variance captured by each principal component after the singular value decomposition (SVD). The first 3 principal components capture 54%, 18% and 10% of the variance, respectively, totalling >80% and thus representing the significant dimensions. **b**. The quality of the representation, cos^2^, for the proteins is plotted for each dimension (small/white: low cos^2^; large/blue: high cos^2^). **c.** The strength (circle size) and direction (blue-positive, red-negative) of the correlation (R^2^) between each protein and the PCA dimensions. **d-f.** The basis vectors for dimensions 1-3 showing the amplitude of each protein in the vector. **g**. Correlations between the plasticity features (columns) identified using the basis vectors (see Results) and PCA dimensions 1-3. Filled cells are significant, Bonferroni corrected correlations (green = positive, red = negative).

#### 2.3.2. Identifying Candidate Plasticity Features

The 3 significant PC vectors are represented by the weighted contribution from each of the 7 proteins that together make up the basis vectors (Fig. 2D-F). Those vectors were used to identify the proteins that drove the variance in the data. That information was stored as XY coordinates in the `pca` object and it was called with the following coding example (6).

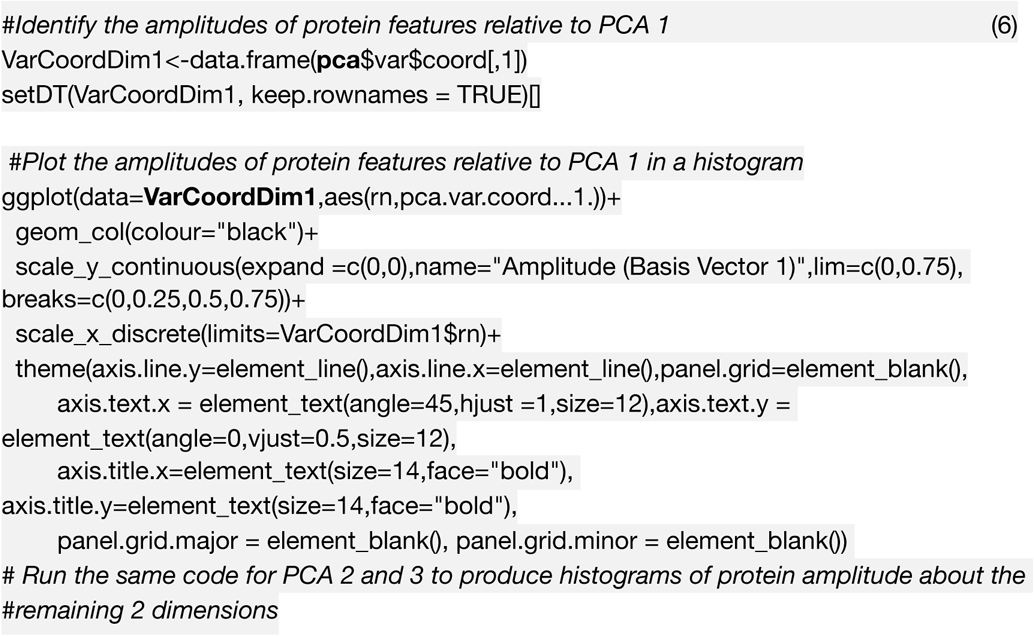

We used a series of steps to identify candidate plasticity features using the output from the PCA, starting with the cos2 metric to assess the quality of the representation for each protein on the dimensions. In the example data set, the glutamatergic proteins GluN1, GluN2A, GluN2B and GluA2 were strongly represented, and GABA_A_α3 was moderately represented on Dim1, and GABA_A_α1 was strongly represented on Dim2. However, synapsin was weakly representation on all of the dimensions (Fig. 2B). Next, we used the correlations between individual proteins and PCA dimensions (Fig. 2C) and the basis vectors (Fig. 2D-F) to find combinations of proteins that would make up the candidate plasticity features. For PC1, we observed that all proteins had positive weights, suggesting that the amount of the proteins was driving that dimension. From PC1, 3 candidate features were calculated: summing all 7 proteins, summing the glutamatergic proteins (GluN1, GluN2A, GluN2B and GluA2), and summing both of the GABAergic proteins (GABA_A_α1 and GABA_A_α3). Next, PC2 and PC3 (Fig. 2E, F) were inspected, and those basis vectors had both positive and negative weights, suggesting that those dimensions represented changes where some proteins increased while others decreased. For example, some pairs of proteins (e.g., GluN2A:GluN2B) that are known to change in opposite directions with different forms of visual experience represented directions. This step also identified novel pairings (e.g., GABA_A_α1:GluN2A) that were included in the set of candidate features. Continuing this approach, we identified 9 candidate features from the 3 basis vectors, and all were combinations of proteins rather than individual proteins. Thus, these steps for using PCA describe an initial dimensionality reduction followed by expansion into candidate plasticity features. Importantly, the expansion steps will identify both novel features and ones that have been well studied, thereby facilitating the interpretation of the results within a biologically relevant framework.

The features were validated by determining the correlation between each of the 9 candidate features and the 3 PC dimensions (Fig. 2G). This was done by calculating the 9 candidate features for all of the samples using the protein expression data and correlating those with the eigenvalues for the 3 dimensions. Bonferroni correction was done to adjust the significance level for the multiple correlations and features that were significantly correlated with a dimension were used as the plasticity features in subsequent stages of the workflow.

The validation of candidate features was done in R, by storing the new features in a matrix `NewFeatures`, centering and scaling those data, then correlating the eigenvalues with the `NewFeatures` matrix. The function *corr.test* from the *psych* package (Revelle, 2019) was used for that step. The strength of the significant correlations was visualized with a custom 2D matrix created using the *geom_tile* function from the *gplots* package (Warnes et al., 2015).

The new features were centered, scaled and stored as a new data matrix:

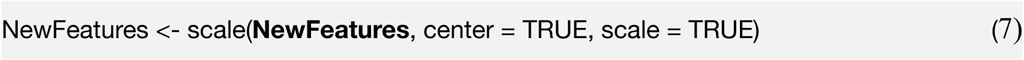

Next, the coordinates for all data points in PCA space were stored in a separate matrix by consulting the `pca` object as follows:

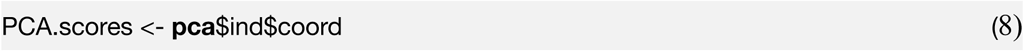

Finally, the correlations between the two matrices `NewFeatures` and `PCA.scores` were determined and visualized using the following R coding example (9):

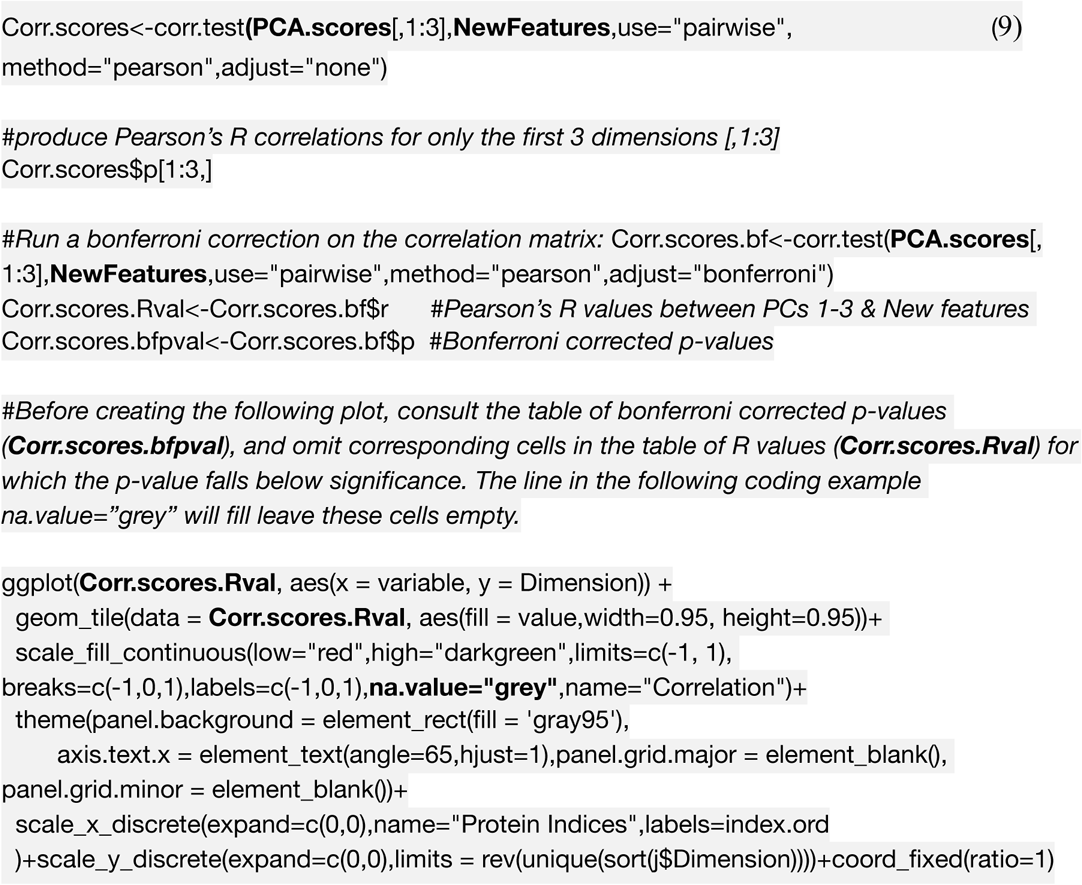

The plot of correlations between the 3 significant PC dimensions and the candidate features was used to validate the suite of features used to construct the plasticity phenotype. In the example above, all of the candidate features except a measure of the E:I balance was significantly correlated with at least one of the dimensions. Interestingly, none of the features were correlated with all 3 dimensions demonstrating that multiple plasticity features are necessary to capture the variance in the data.

#### 2.3.3. Using the Plasticity Features to Construct a Plasticity Phenotype

The collection of plasticity features were combined to form the *plasticity phenotype* that was visualized using the following code (coding sections 10 & 11).

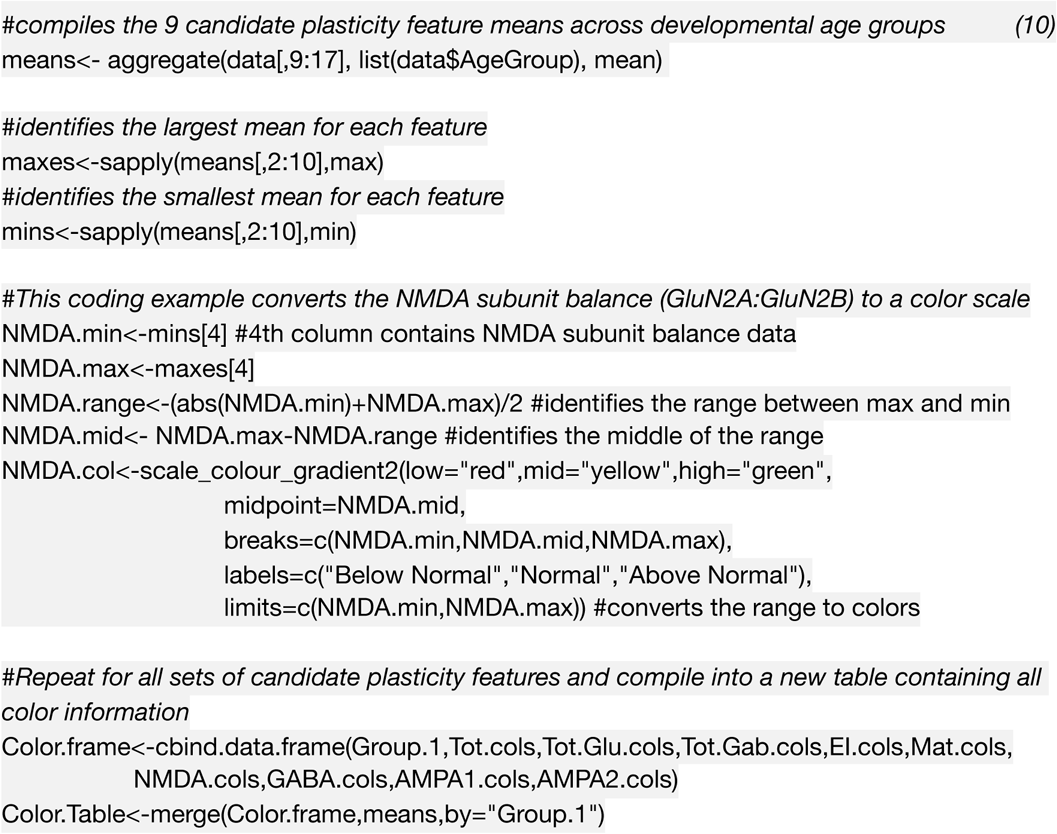

That list of colour-codes for each subcluster was stored in a new matrix called `Colour.Table`. The matrix will be consulted in the code below (11) to call the correct colour for each horizontal bar in the plasticity phenotype.

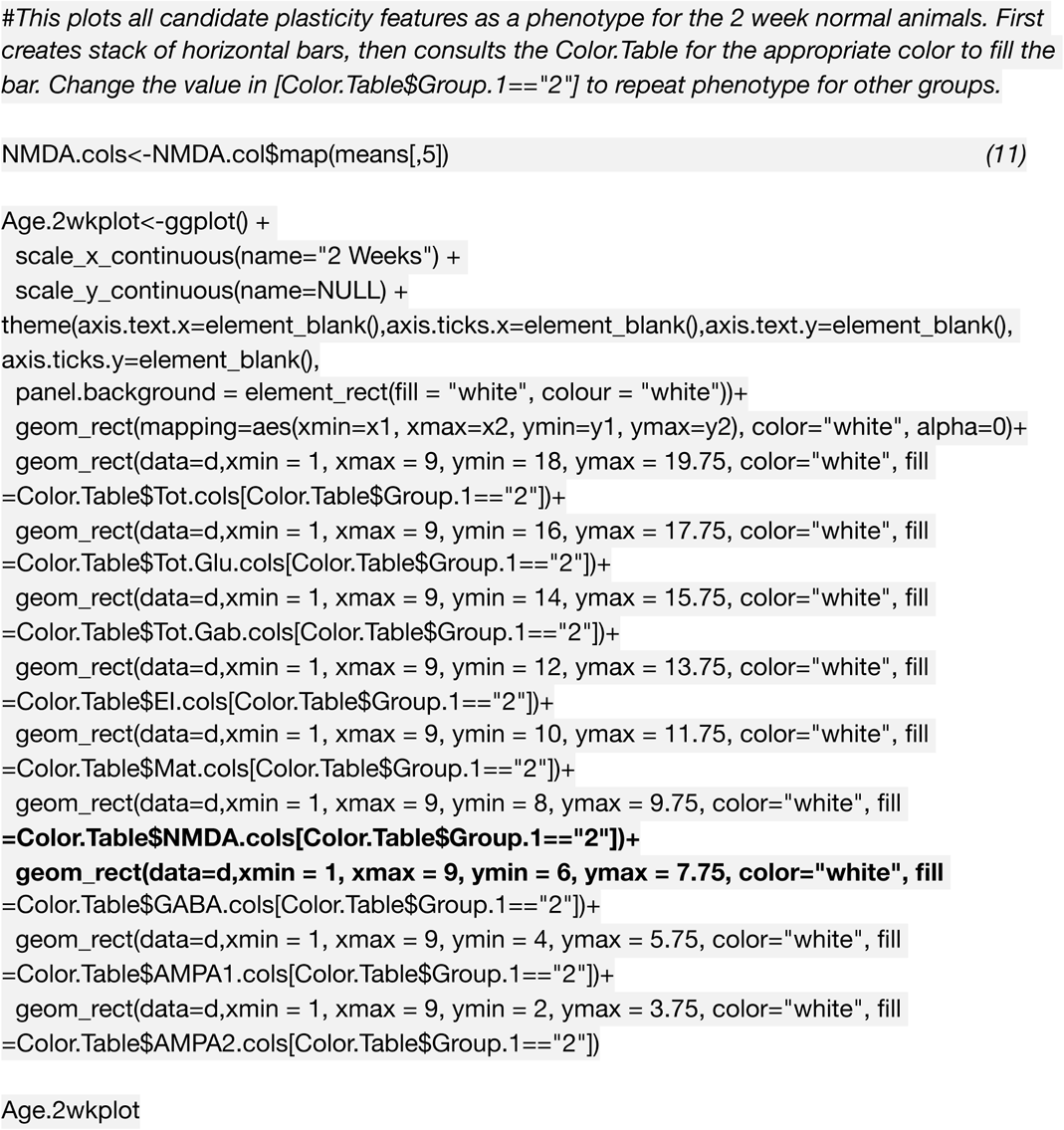

#### 2.3.4. Interpretation of the plasticity phenotype for studying V1 development

In the example, the development of each plasticity feature is calculated and displayed as the average plasticity phenotype for V1 at each of the ages studied (Fig. 3A). The colour-coded stack of 9 bars provides a compelling visualization of the plasticity phenotypes that aids in seeing the pattern of developmental changes in V1. The top 3 grey-level bars highlight steady increases in the amount of protein expression up to 12 weeks before declining to adult levels at 16 weeks of age. Also, the visualization of the plasticity phenotype helped to see patterns that were particular to different age groups. In the younger ages, the 6 indices had similar patterns, but by 8-12 weeks of age, the pattern was different, and at 16 weeks of age, it appeared similar to the adult. Thus, using all 9 features in the plasticity phenotypes helped to identify the peak of the critical period in cat V1 from 3-6 weeks and the end at 16 weeks. The boxplot for each feature allows for additional inspection of the differences and statistical comparisons among the ages (Fig. 3B-J). We have found that the plasticity phenotype is particularly for finding protein indices that are not typically studied. An example is the GABA_A_α1-GluN2A index, which was strongly correlated with PC Dim 2. That index shifts from more GABA_A_α1 to more GluN2A during the development of V1. Although the expression of both GABA_A_α1 and GluN2A increases during maturation (Beston et al., 2010; Bosman et al., 2005; Sheng et al., 1994) the shift supports the idea that GABAergic processes dominate during the critical period and glutamatergic processes in adulthood.

**Figure 3.**
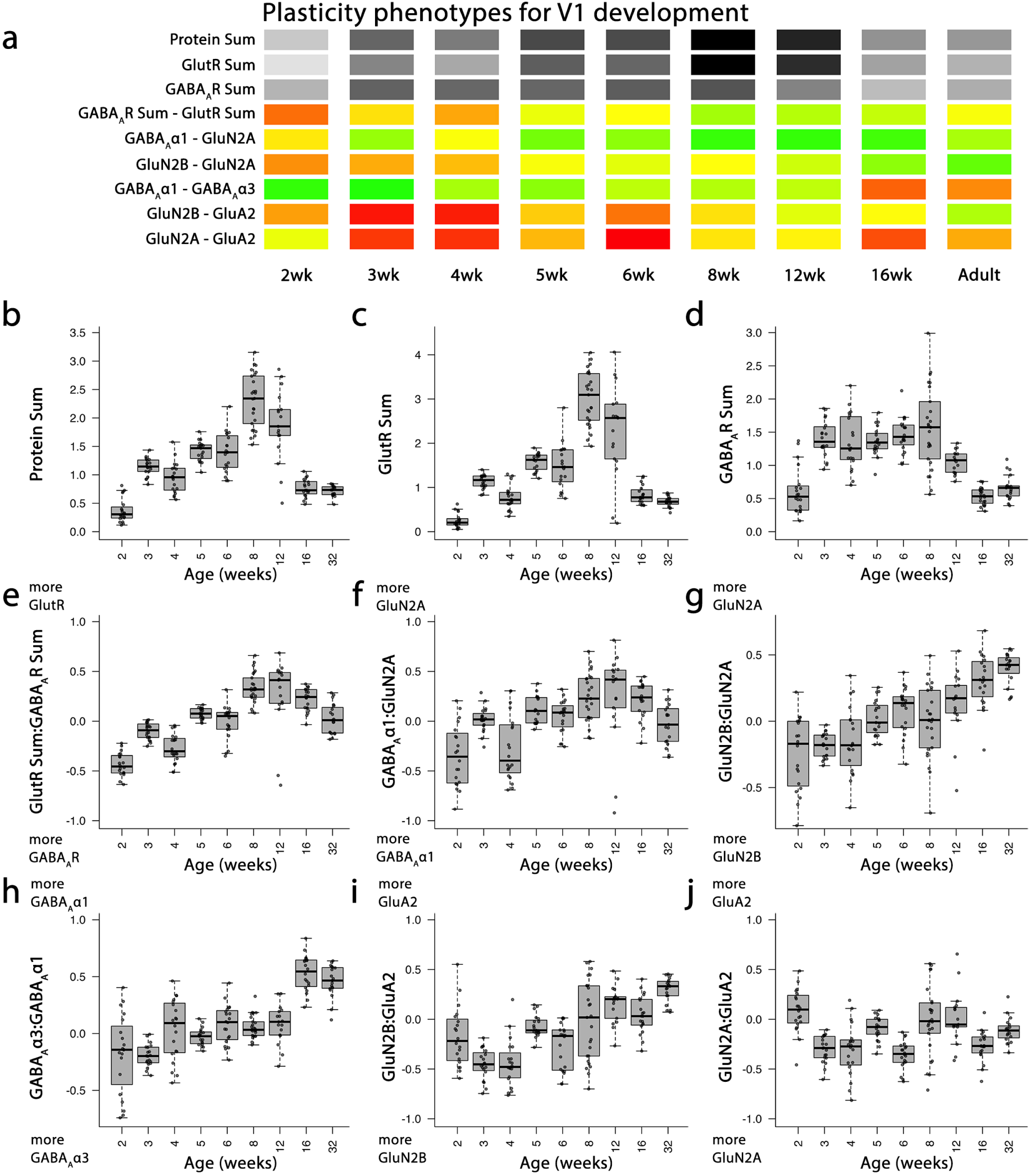
Plasticity phenotypes for V1 development. **a**. The developmental sequence of plasticity phenotypes in cat V1 was visualized as a stack of colour-coded horizontal bars. The 3 grey-level barsat the top represent the protein sums and the 6 red-green colour-coded bars represent the protein indices identified by the PCA. The plasticity phenotypes were calculated for each stage of development and ordered from youngest to oldest. **b-j** The boxplots show the average protein sum **(b-d)** and an average index value **(e-j)** for each of the stages of development.

### 2.4. Application of Plasticity Features and Phenotypes for Clustering of Experience-Dependent Changes in V1

In this section, we describe a workflow that uses the plasticity features and phenotypes as the input to a clustering algorithm for partitioning the data into clusters and subclusters (Fig. 4). This approach is useful when asking questions about changes in V1 after different types of visual experience where the goal is to assess the similarities and differences among rearing conditions. Also, this workflow is helpful for discovering novel associations in the data. Here we illustrate the steps using protein expression data from V1 of animals that had different types of visual manipulations (RO, BD or BV) to promote recovery from early monocular deprivation (MD) (Balsor et al., 2019).

**Figure 4.**
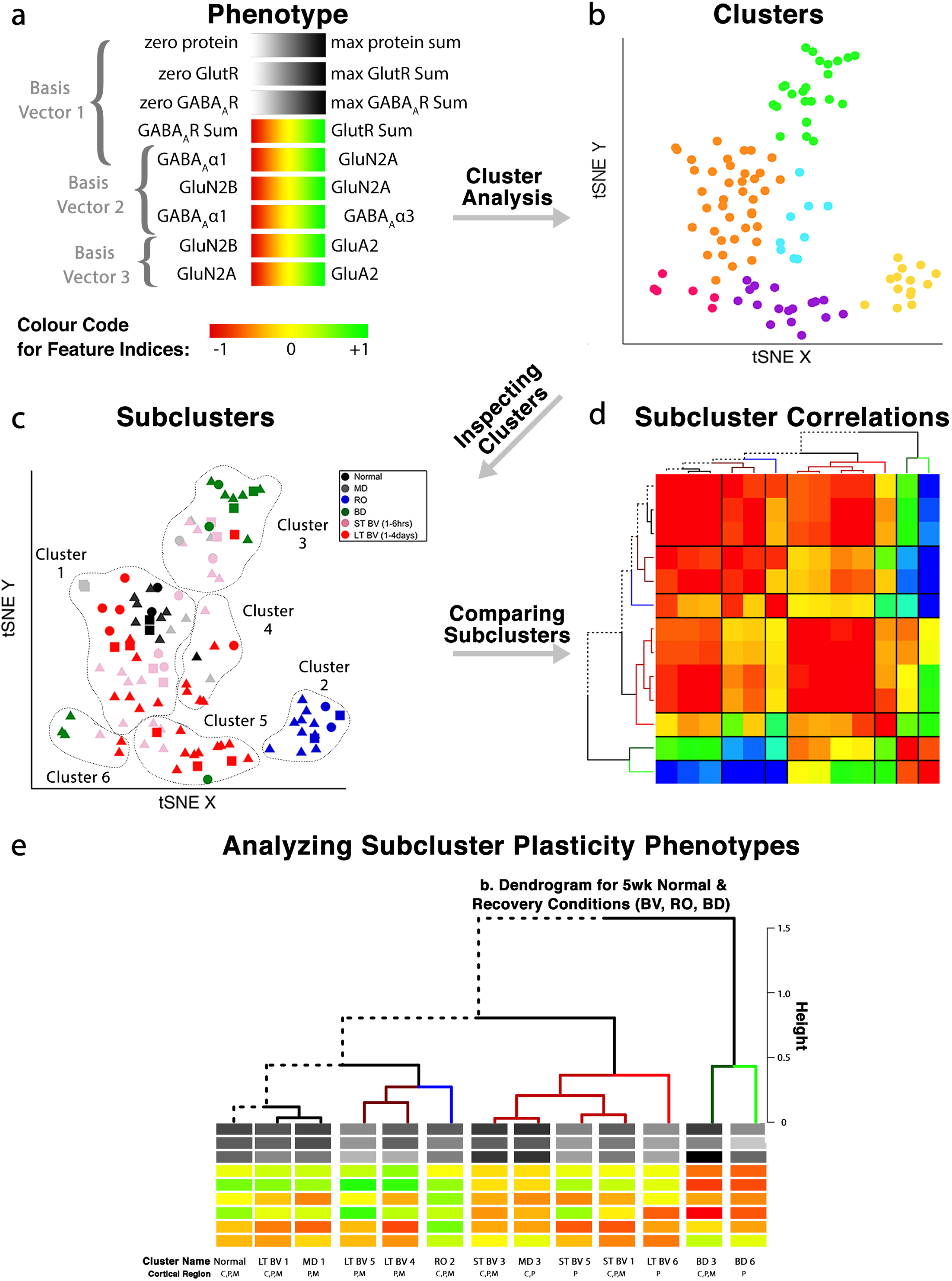
Analysis workflow to use plasticity phenotypes in cluster analysis. The analysis workflow is shown for using the plasticity phenotypes to identify clusters and subclusters. **a**. The legend for the plasticity phenotype where each feature is represented as a series of stacked, horizontal bars, and the top 3 grey-level bars and bottom 6 coloured bars represent protein sums and indexes, respectively. **b, c.** The 9 plasticity features were used as the input for the tSNE analysis, and k-means clustering was applied to the transformed data. **b**. The tSNE identified clusters are outlined. **c**. The composition of the clusters was inspected by coding the rearing conditions using different coloured dots. **d**. Samples from the same rearing condition and cluster are divided into subclusters, and the strength of the pairwise correlations among the plasticity phenotypes is shown using a correlation matrix. **e**. The plasticity phenotypes for each subcluster were displayed at the ends of the dendrogram that ordered the subclusters (**d**).

This application of the plasticity features and phenotypes uses tSNE analysis to partition the data into clusters. tSNE preserves both the global and local arrangement of the plasticity features and is a good way to visualize clusters because it artificially scales the distance between data points with similar patterns of features. There are, however, many other clustering methods that could be used for this step and the selection of the most appropriate clustering algorithm will depend on the structure of the data.

In this example, the inputs to tSNE are the validated plasticity features (`NewFeatures`) without identifying information such as the cortical region, rearing condition, or age.

The following coding example (12) demonstrates how to perform a tSNE analysis using the *tsne* function from the *tsne* package (Donaldson and Donaldson, 2010).

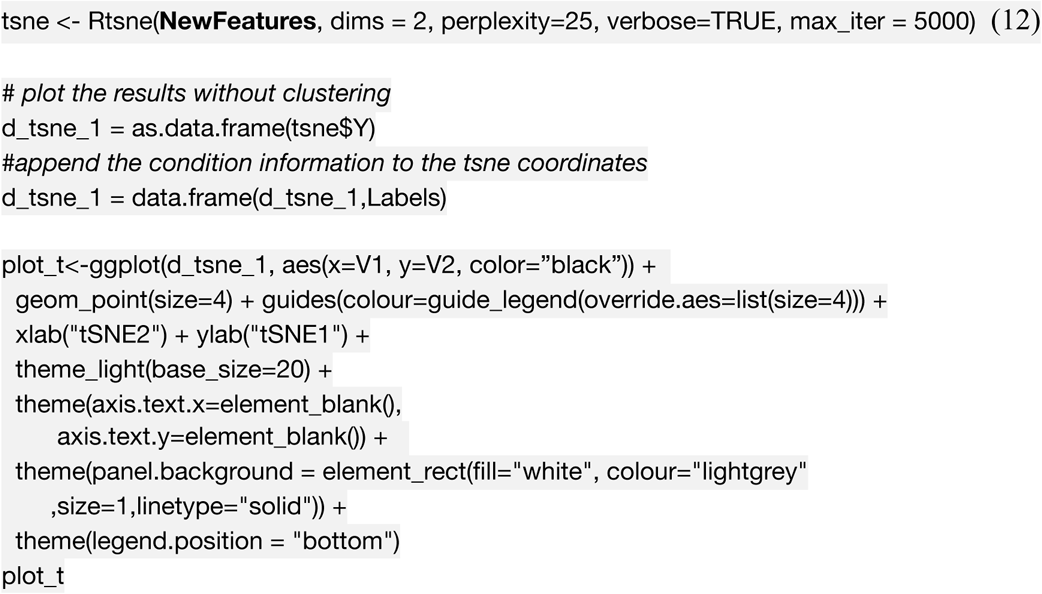

The first step in the tSNE analysis reduced the plasticity features from each sample to a 2-dimensional set of tSNE XY coordinates (Fig. 5A). Those coordinates were used as the input to *K*-means clustering analysis to identify and then visualize clusters in the data set.

**Figure 5.**
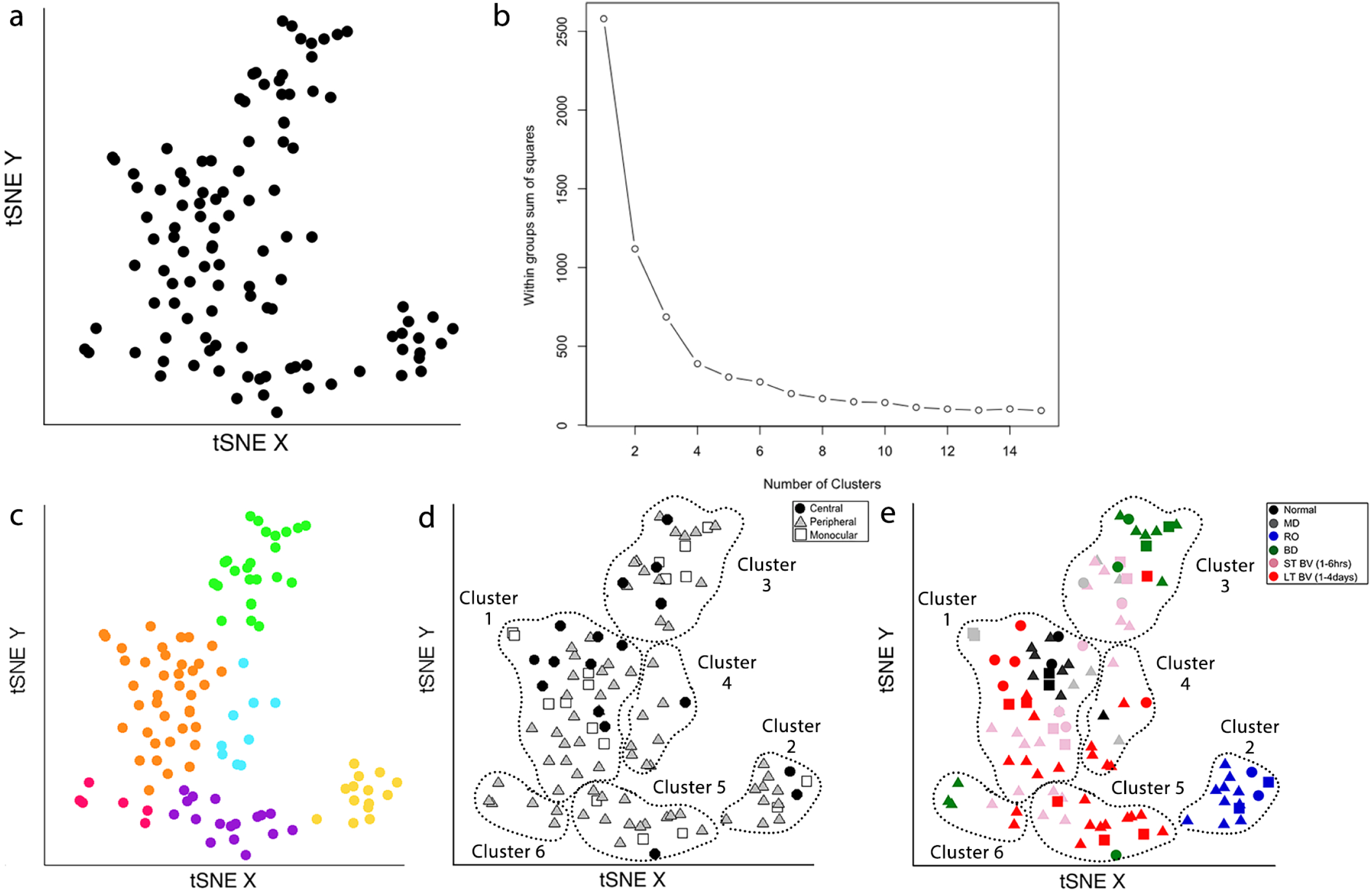
Identifying clusters from plasticity phenotypes using t-Distributed Stochastic Neighbor Embedding (t-SNE) and k-means clustering. **a**. An XY representation of tSNE transformed data. **b**. The optimal number of clusters was chosen by measuring the within-groups sum of squares, and choosing the inflection point (k=6). **c**. The 6 clusters are represented as different coloured dots. **d**. The content of each cluster was visualized for the 3 cortical regions (central, peripheral, monocular) and **e**. 6 rearing conditions (normal, MD, RO, BD, ST-BV, LT-BV).

Both *K*-means and hierarchical clustering algorithms require the number of clusters *k* as a parameter. A good method for choosing the number of clusters is to measure the within groups sum of squares (WSS) for a range of *k*, plot that information and then determine the inflection point.

In the example data set there were 9 rearing conditions (e.g. normal, monocular deprivation etc) so we chose a range for *k* of 2 to 15 clusters, that encompassed the number of conditions.

The following coding example (13) determined the WSS and plotted it for the *k* clusters:

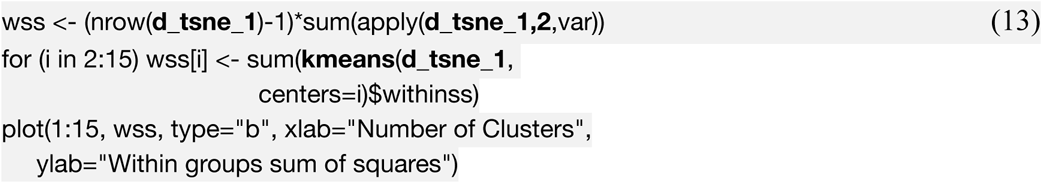

The optimal number of clusters was selected by fitting an exponential decay curve to the WSS data then finding the number of clusters corresponding to the point where the curve plateaued (4τ) (*k*=6). This approach is called the “elbow method”, where 4τ is the point of inflection, or elbow, of the curve (Fig. 5B).

Next, *K*-means clustering for *k*=6 was done on the output from the tSNE analysis (Fig. 5C). The clusters were assigned different colours to visualize the samples in each cluster. Some clusters (green and yellow) were spatially separated on the tSNE plot, while others (e.g., orange and blue) were adjacent. The following coding example (14) plots the clusters identified in the tSNE representation of the data as different colors, but these colors can be manipulated to represent other characteristics of the data (e.g. cortical area, treatment condition).

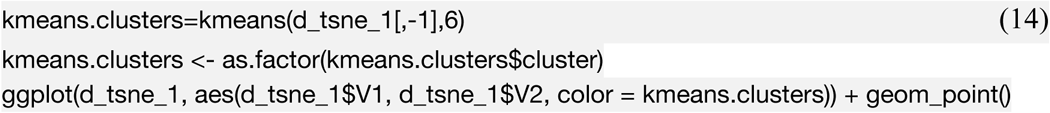

The number of samples in each cluster ranged from 5 (magenta) to 38 samples (orange). We annotated each sample based on the visual cortical region (central, peripheral, or monocular) (Fig. 5D) and rearing condition (Fig. 5E) to analyze cluster composition and determine if the clustering reflected one of those parameters. For example, cluster 2 contained samples from only one rearing condition (reverse occlusion) and cluster 1 contained almost all of the normally reared cases but it also had samples from other rearing conditions. Thus, this step identified clusters and provided some evidence that the rearing condition was driving changes in the plasticity phenotypes. The tSNE clustering, however, did not reveal which features from the phenotypes were separating the samples into different clusters or grouping them into the same cluster.

Annotating each sample by the cluster, visual cortical region, and rearing condition was an essential first step for using the plasticity phenotypes to identify and explore *subclusters* in the data. That process identified 13 subclusters in the example data set (Table 3).

**Table 3.** Subclusters identified from the tSNE plot.

| Subcluster | Rearing Condition | tSNE Cluster | Region |  |  |
| --- | --- | --- | --- | --- | --- |
|  |  |  | C | P | M |
| Normal 1<sub>C,P,M</sub> | Normal | 1 | X | X | X |
| LTBV1<sub>C,P,M</sub> | Long term BV recovery | 1 | X | X | X |
| MD1<sub>P,M</sub> | Monocular deprivation | 1 |  | X | X |
| STBV1<sub>C,P,M</sub> | Short term BV recovery | 1 | X | X | X |
| RO2<sub>C,P,M</sub> | Reverse occlusion | 2 | X | X | X |
| STBV3<sub>C,P,M</sub> | Short term BV recovery | 3 | X | X | X |
| MD3<sub>C,P</sub> | Monocular deprivation | 3 | X | X |  |
| BD3<sub>C,P,M</sub> | Binocular deprivation | 3 | X | X | X |
| LTBV4<sub>P,M</sub> | Long term BV recovery | 4 |  | X | X |
| LTBV5<sub>P,M</sub> | Long term BV recovery | 5 |  | X | X |
| STBV5<sub>P</sub> | Short term BV recovery | 5 |  | X |  |
| LTBV6<sub>P</sub> | Long term BV recovery | 6 |  | X |  |
| BD6<sub>P</sub> | Binocular deprivation | 6 |  | X |  |

A final note: In this workflow, dimensionality reduction and feature selection were performed before tSNE analysis and clustering. Although this is a common approach for analyzing high-dimensional data in neuroscience it is important to remember that PCA preserves features with variance that is aligned with the orthogonal dimensions. Thus, features with more subtle but important variance away from the PCA dimensions will not be included in subsequent clustering (Chang, 1983).

### 2.5. Identifying and Exploring Subclusters using Plasticity Features and Phenotypes

Depending on the design of the study there may be additional factors that lead to subclustering of the data. In this section, we describe a method to analyze and visualize subclusters using the features that comprise the plasticity phenotypes.

First, the features and tSNE results were combined in R by appending the object containing the tSNE dimensions and clusters (`d_tsne_1`) to the plasticity features (`NewFeatures`), and stored in a data matrix (`data`). Now each sample had both the clustering information from the tSNE analysis and the feature data from PCA. Next, the data were organized into subsets according to the subclusters. For example, all of the data points for Normal samples in cluster 1 were subset as follows:

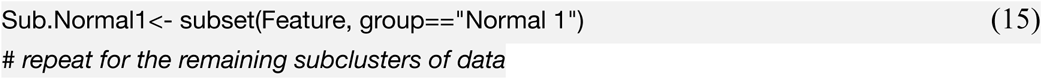

It is a daunting task to try and synthesize all of the significant differences for 9 features and 13 subclusters. To facilitate this we calculated the pairwise correlations between subclusters for the plasticity phenotypes, ordered the subclusters using hierarchical clustering and visualized the correlations in a 2D heatmap (Fig. 6). The steps are described below in coding example 16.

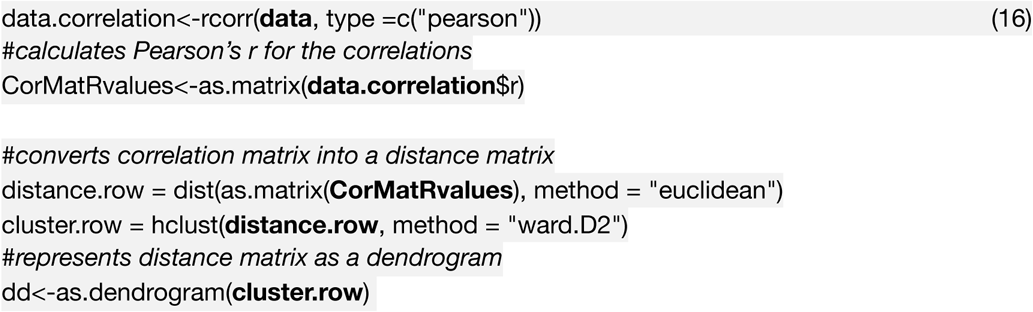

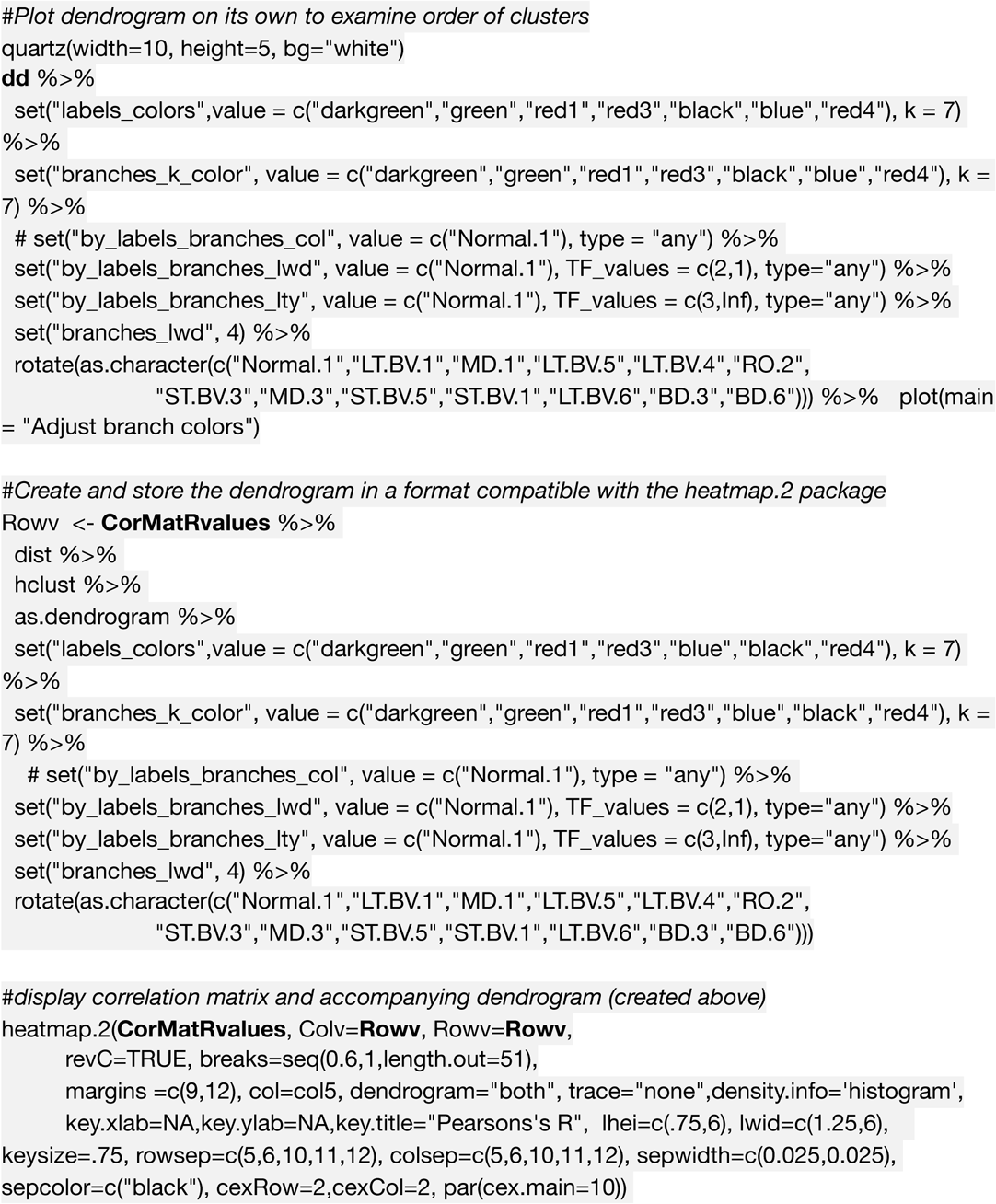

**Figure 6.**
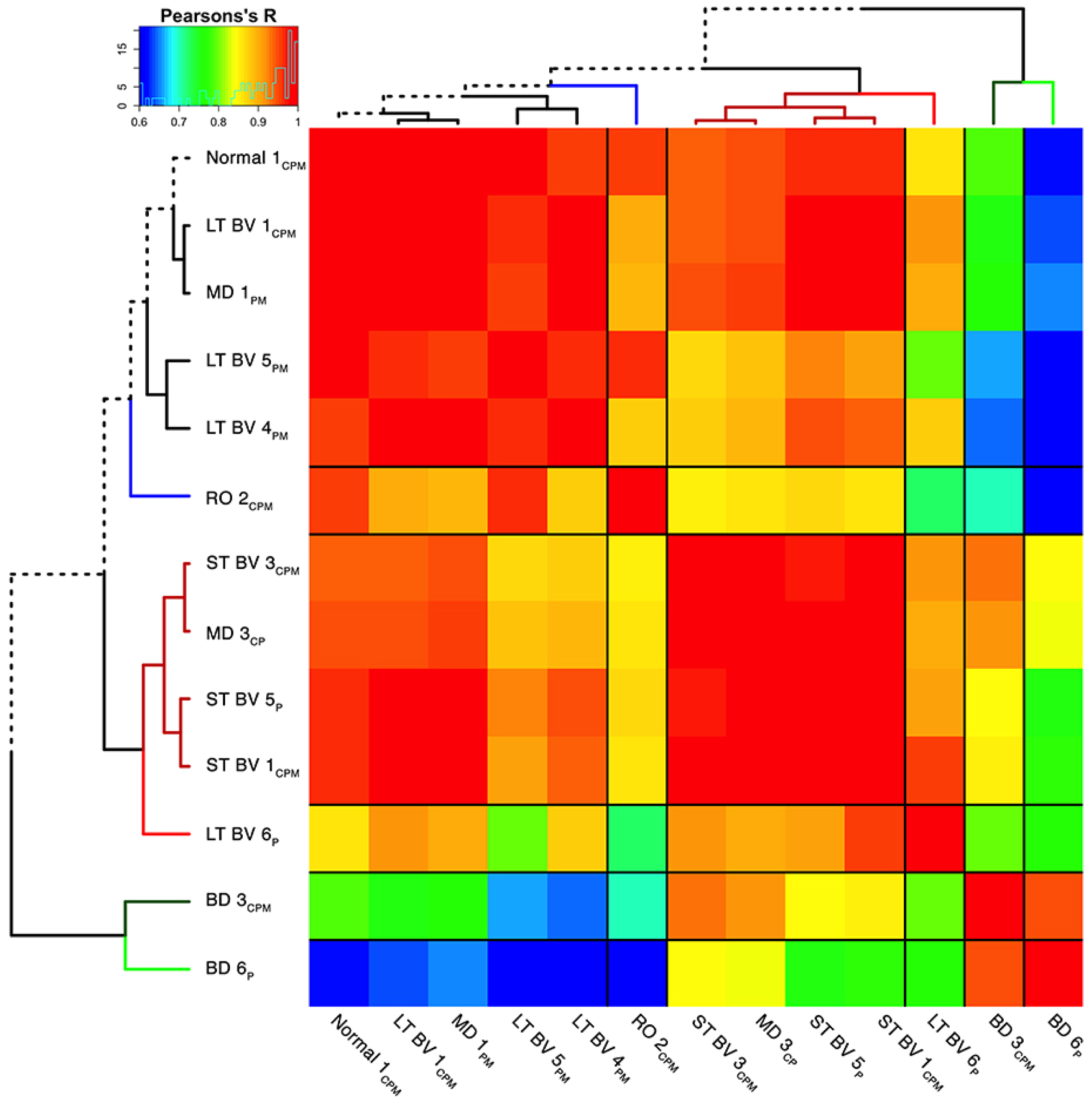
Visualizing pairwise correlations between treatment subclusters. The matrix shows the strength of correlation (colour) between the pairs of subclusters identified in the previous step. Subclusters were ordered using hierarchical clustering to position subclusters with stronger correlations nearby in the matrix. This reordering identified 5 groups based on the height of the dendrogram branches and marked by different coloured lines in the surrounding dendrogram. The solid black lines in the matrix mark the 5 groupings of subclusters.

The correlation matrix for the plasticity phenotypes showed the strength of similarity or dissimilarity among the subclusters (Fig. 6). Here, the surrounding dendrogram ordered subclusters for some rearing conditions (e.g., LT BV) on the same branch as the Normal subcluster, while other conditions (e.g., BD) were far from the Normal branch. This analysis showed which subclusters had similar plasticity phenotypes but did not clarify if the similarity was based on the entire pattern of the plasticity phenotype or if a smaller number of features drove the clustering.

### 2.6. Constructing and Visualizing the Plasticity Phenotypes for the Subclusters

In the last step for this workflow, we describe visualizing the plasticity phenotypes, ordering them using the dendrogram from the hierarchical clustering (Fig. 7), and comparing phenotypes to identify differences among rearing conditions.

**Figure 7.**
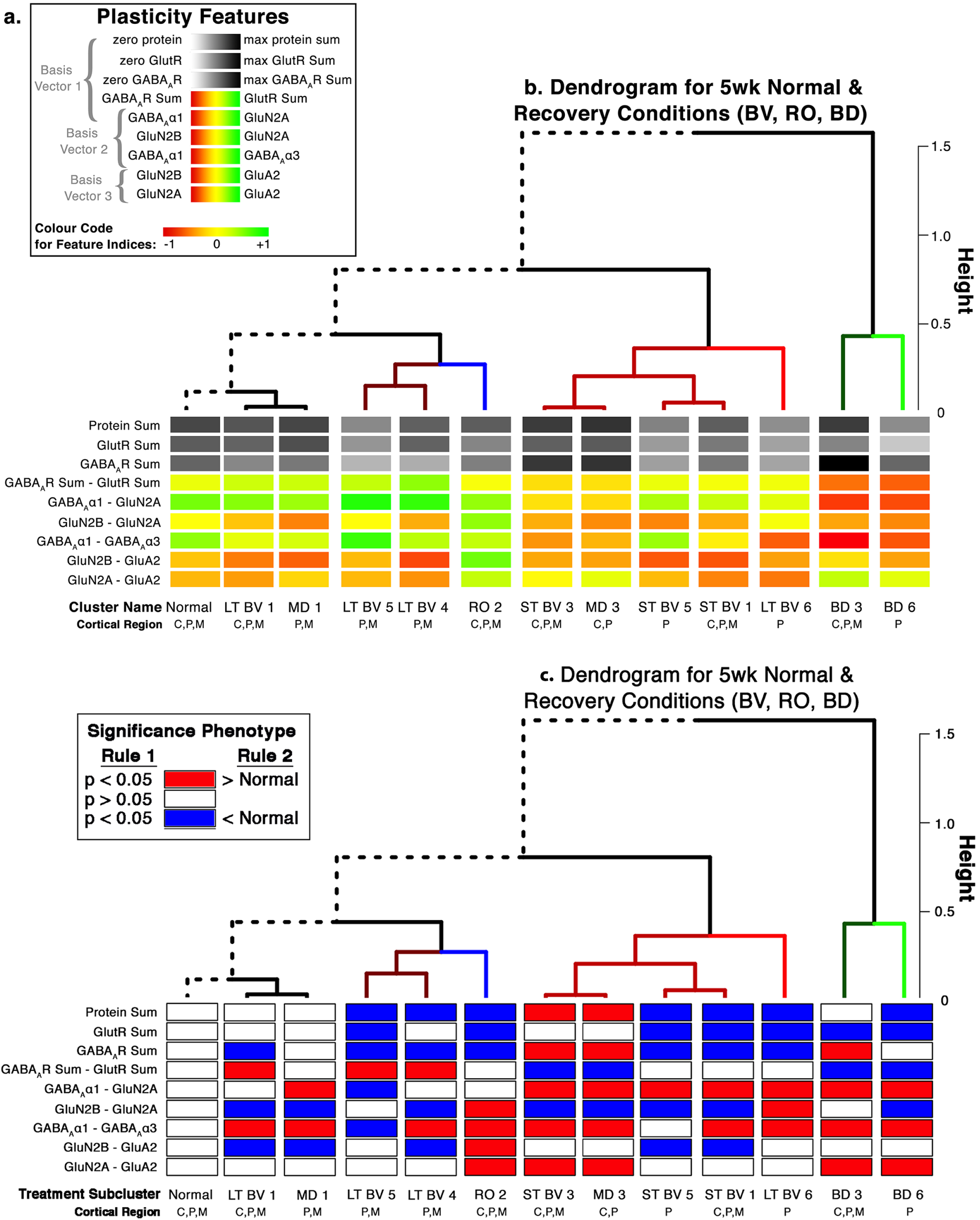
Analysis of plasticity phenotypes for the different rearing conditions and subclusters. **a.** The plasticity phenotype legend with the same conventions as Figure 4a. **b.** The plasticity phenotypes for each subcluster were ordered using the dendrogram that surrounds the correlation matrix in Fig. 6. **c.** The plasticity features for treatment subclusters that differed from the normal subcluster were visualized using a layout that paralleled the design of panel **b**. Significant differences were identified by colour-coding the plasticity feature band red if the feature was > normal and blue if it was < normal (p<0.05). The colour convention is shown in the figure inset.

A display was created to show each of the feature and the whole pattern of the plasticity phenotype so that it was easy to inspect the subclusters for similarities and differences visually (Fig. 7A). The plasticity phenotypes were visualized in R, using the *geom_tile* function in the *ggplot2* package (Wickham, 2009). First, the feature mean was determined for each subcluster then the limits of the colour scales were set by finding the maximum and minimum expressions for a feature across all subclusters. Finally, the subcluster mean was converted to the corresponding RGB score. The following coding examples (17 & 18) was used to map the mean for each feature in the Normal condition onto a colour scale:

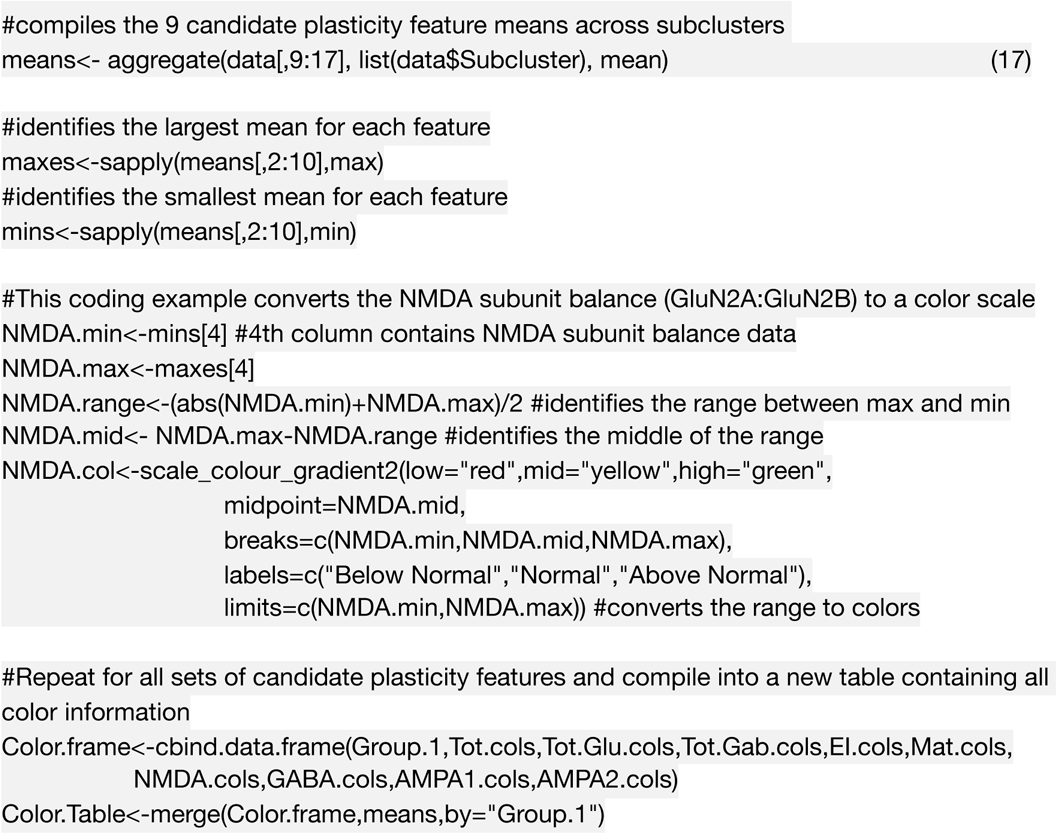

That list of colour-codes for each subcluster was stored in a new matrix called `Colour.Table`. The matrix will be consulted in the code below (18) to call the correct colour for each horizontal bar in the plasticity phenotype.

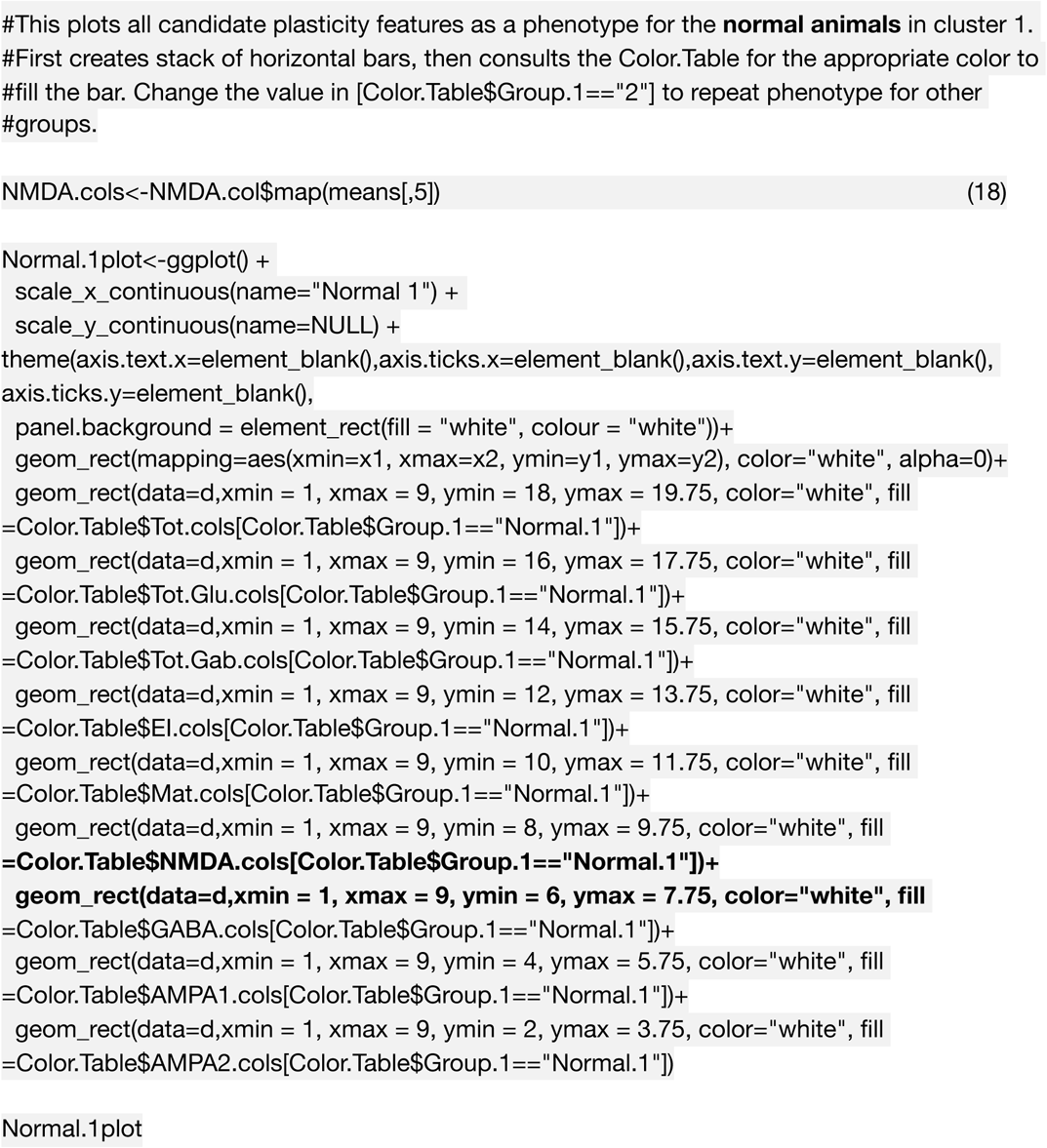

#### 2.6.1. Interpretation of the plasticity phenotype and cluster analysis for classifying experience-dependent changes in V1

Figure 7 illustrates the plasticity phenotypes for each of the subclusters and the surrounding dendrogram arranges them from the phenotypes that are most similar to normal on the left side to progressively dissimilar moving to the right (Balsor et al., 2019). Using this visual tool helps to see which features contribute to partitioning the rearing conditions into different subclusters. For example, although reverse occlusion (RO) and binocular deprivation (BD) conditions cause poor acuity in both eyes it is easy to see that their plasticity phenotypes are different and the features have shifted away from normal in different directions (Fig. 7A). Further inspection of the phenotypes identified other differences in the pattern of features among the subclusters. In addition, the same layout of the phenotypes can be used to identify which features are different from normal. In this example, we have colour-coded the feature bars red for above and blue for below normal levels (Fig 7B).

The subclusters in this example were ordered using the same hierarchical clustering dendrogram as in Figure 6, and the average plasticity phenotype for a subcluster was displayed at the end of its branch in the tree (Fig. 7B). The figure provides a strong visual impression of the phenotypic similarity among subclusters located nearby in the tree (e.g., Normal and LT BV) and differences for subclusters that are further away (e.g., Normal and BD). Thus, this tool supports linking the output of high-dimensional analyses with neurobiologically meaningful insights.

Finally, features that were significantly different from Normal were visualized using a graphic similar to the phenotype tool except that the bars were colour-coded red if the feature was above- or blue if the feature was below-Normal (Fig. 7C). The statistical comparisons were done using a Bootstrap analysis described in the Supplemental Analysis section. Features that did not differ from Normal were left empty in the figure. This method is an effective way to visualize and present a large number of statistical comparisons among the features.

Identifying features that were significantly different from normal facilitated attaching biological meaning and classifying V1 changes driven by the rearing conditions. For example, the top 3 features of the RO subcluster were significantly less than normal indicating generalized loss of protein expression, while the bottom 4 features were significantly greater than normal suggesting abnormal protein balances favouring GluN2B and GABA_A_α1. A notable finding from this analysis was that even though the BV treatment conditions clustered near the normals, none of the treatments returned all of the plasticity features to normal levels.

## 3. Discussion

Many neural proteins underpin the complex set of processes that support experience-dependent plasticity in V1. In this paper, we have provided a step-by-step primer using R code to illustrate a data-driven approach for combining measurements from multiple plasticity proteins to make and use a plasticity phenotype for studying V1. The examples highlight the steps for discovering the suite of proteins that make up plasticity features and then how to combine those features into a phenotype. We illustrated the use of the plasticity phenotype to classify tissue-level changes in V1 for different ages and rearing conditions. Also, we showed that the visualization tool for the plasticity phenotype helps synthesize large amounts of data into a single figure for exploring V1 development and changes after visual experience. This application of a phenotype has the potential to guide discoveries about new relationships between experience-dependent driven development of V1 and vision.

In this paper, we have used the term “phenotype” to classify a suite of plasticity-related proteins that are known to interact with the visual environment. Thus, the *plasticity phenotype* can be thought of as a tool for classifying a set of plasticity mechanisms in V1 that support the development of vision and ongoing dynamics specific to each individual’s variations in perceptual experience. The workflows described in the paper were developed to help with interpreting complex data sets that include the expression of multiple proteins or genes at different ages or rearing conditions. Aspects of the workflows are similar to other approaches used to study V1 that applied sequential steps beginning with dimension reduction (e.g. PCA, tSNE) and then cluster analysis (e.g. (Carlyle et al., 2017; Luo et al., 2017)). The new workflows extend previous work by developing steps to build, visualize, and compare plasticity phenotypes to capture neurobiological features that characterize developmental stages or types of visual experience. The plasticity phenotype represents a new heuristic for analyzing V1 development and plasticity that can facilitate meaningful interpretation for intricate patterns of proteins or genes.

Since the current workflows build on established methods for high-dimensional data analysis, it is possible to adapt the code to use other algorithms by exchanging a few lines in the R code. New tools such as pcaExplorer may also help by providing an interactive interface for using PCA to explore and interpret gene or protein expression patterns (Marini and Binder, 2019). Also, as new plasticity mechanisms are identified, those can be explored to find novel associations between visual plasticity and V1 neurobiology. The advent of single-cell transcriptomics using RNA sequencing to study brain development has added another layer of complexity but those very high-dimensional data studies have spurred the development of new analytical tools to discover gene markers for cell phenotypes (Aevermann et al., 2018). We are just beginning to appreciate the complexity of these associations. There is little doubt that further development of high dimensional analyses will be necessary for decomposing the direct and indirect effects of changing patterns of plasticity proteins and genes on visual development. Below we discuss some of the natural next steps in methodology development.

Using the plasticity phenotype worked well with the example data sets, and it facilitated classifying patterns of changes in V1 during development and after abnormal visual experience, but this approach needs to be tested with other data sets. In particular, the feature selection steps (Section 2.3.2) were supervised. We took advantage of prior knowledge about the function of the plasticity proteins to select suites of proteins to comprise the features. Additional work is needed to develop unsupervised methods for identifying features and transforming data. These changes will be especially important when working with very high dimensional data sets containing hundreds or thousands of proteins or genes and looking for new suites of plasticity features that may have a role in V1 development. Here, other methods for solving the feature selection problem, such as minimum redundancy maximum relevance (mRMR) (Ding and Peng, 2005), or random forest machine learning (Aevermann et al., 2018) may be helpful for automatic annotation of patterns in the data.

Similarly, an exploratory process and manual inspection of clusters were used in the clustering workflow to select the k-means parameters for the number of clusters (Section 2.4, coding example (13)). Estimating the number of clusters in a data set is challenging; however, methods such as the gap statistic can be added to the workflow for choosing the number of clusters (Tibshirani et al., 2001). Additionally, recent approaches to clustering, such as robust (weighted) sparse k-mean clustering, have the advantage of simultaneously identifying clusters and informative features for partitioning the data that can be used in feature selection (Brodinová et al., 2019). Finally, growth mixture models for cluster analysis of longitudinal data may be more suitable for data analysis from studies that include a series of sequential measurements of cortical development (Wei et al., 2017).

The development of high-throughput methods for measuring large numbers of proteins and genes are changing how plasticity is studied in V1 and other cortical areas. These large data sets are enabling a deeper understanding of the subtle changes in V1 driven by visual experience and a richer appreciation of individual variations. In other areas of biomedical research, the use of phenotypes has accelerated the discovery of associations between human traits and genetic variants. Furthermore, the term plasticity phenotype has been used as a description for the waxing and waning of plasticity-related gene expression in V1 (Smith et al., 2019). In this primer, we have described an extension of the phenotype heuristic to represent suites of plasticity-related proteins that change during development or after abnormal visual experience. Our goal in constructing a plasticity phenotype was to facilitate the translation of complex high dimensional patterns of protein or gene expression in V1 into a simpler form that aids in the identification of neurobiological changes affecting experience-dependent development of vision. We have provided code for constructing and using the plasticity phenotype, as well as illustrated some insights about V1 plasticity that our studies have found using this approach (Balsor et al., 2019). Identifying all of the neurobiological mechanisms that underpin the development and plasticity of complex visual traits is still a distant goal. However, the development and use of heuristics such as the plasticity phenotype described in this paper will be necessary tools for studying complex high dimensional data sets of plasticity proteins or genes. Ultimately, the development of phenotypes that cross levels of analysis from molecules to cells to circuits will lead to a deeper understanding of the associations between V1 neurobiology and visual-traits.

We have posted an R package “v1hdexplorer” that aggregates the various packages and custom visualization code used in this paper, and is available for download using the function install_github(“balsorjl/ v1hdexplorer”). The custom visualization scripts are included in this document (e.g. coding examples (10 & 11) and (17 & 18) for visualizing the plasticity phenotypes).

## 5. Supplemental Analysis

### 5.1. Bootstrap analysis using custom R code

To examine the probability that observations in one group (e.g. reverse occlusion-RO) were statistically different from another group (e.g. Normal animals) we performed a one-way bootstrap analysis. First, the *observed* parameters of the experimental group (RO) were used to create a *simulated* data set with 1,000,000 points that had the same mean (`mean.RO`) and standard deviation (`stdev.RO`) as the observed subset of RO data. The object `sim.RO` contains the simulated data set of RO samples.

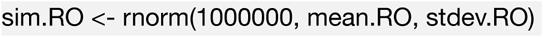

Next, we modelled *experimental* data of the normal group by drawing, at random, from the *simulated* RO data set (`sim.RO`) the same number of samples as the *observed* subset of Normal animals (`NNormal`). We calculated the mean of this first *experimental* subset of data, and replaced all data points in `sim.RO`. This experiment was repeated 100,000 times, and stored in the object `resamples.NormalRO`.

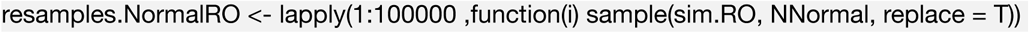

Each of these 100,000 experiments were reduced to the experimental mean. This subset of data represent a simulated population of comparisons between Normal and RO animals. We calculated the mean of this simulated population, and saved it as `mean.resamples.NormalRO`.

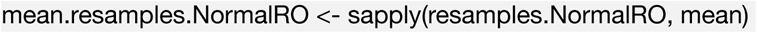

To determine the probability that the mean of the *observed* Normal subset of data (`mean.Normal`) came from the observed RO subset of data (`mean.RO`), we compared `mean.Normal` against the population in `mean.resamples.NormalRO`. To calculate the exact probability (p-value) that the observed Normal group (`mean.Normal`) came from the `mean.resamples.NormalRO` population, we calculated the percentage of data points that fall above or below `mean.Normal` in the distribution of data points in `mean.resamples.NormalRO`.

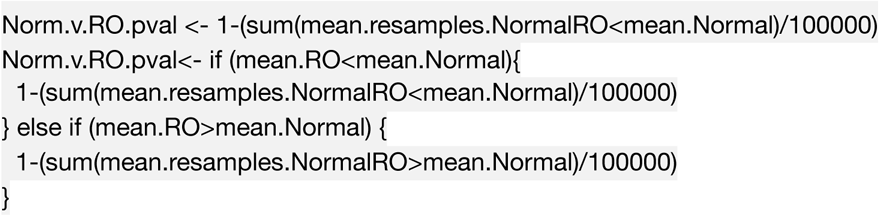

In order to compare other conditions against the normal subset of data, adjust the subset of data that is called in place of RO (eg. BD versus Normal). In order to change the direction of the comparison against Normal to be against a different group (eg. MD), exchange the comparator group for Normal in all of the above statements (eg. BD versus MD). The resultant data can be called into a text file and stored for later use when constructing histograms.

## Acknowledgements

We thank Ewalina Jeyanesan, Keon Arbabi and Pedro Ballester for helpful comments on the manuscript, and Drs. Beston, Pinto, Williams, Beshara and Siu for sharing data used to develop the workflows.

## References

Aevermann, B. D., Novotny, M., Bakken, T., Miller, J. A., Diehl, A. D., Osumi-Sutherland, D., et al. (2018). Cell type discovery using single-cell transcriptomics: implications for ontological representation. Hum. Mol. Genet. 27, R40–R47. doi:10.1093/hmg/ddy100.

Balsor, J. L., Jones, D. G., and Murphy, K. M. (2019). Classification of Visual Cortex Plasticity Phenotypes following Treatment for Amblyopia. Neural Plasticity 2019, 2564018–23. doi:10.1155/2019/2564018.

Benoit, J., Ayoub, A. E., and Rakic, P. (2015). Transcriptomics of critical period of visual cortical plasticity in mice. Proceedings of the National Academy of Sciences 112, 8094–8099. doi:10.1073/pnas.1509323112.

Beston, B. R., Jones, D. G., and Murphy, K. M. (2010). Experience-dependent changes in excitatory and inhibitory receptor subunit expression in visual cortex. Front Synaptic Neurosci 2, 138. doi:10.3389/fnsyn.2010.00138.

Bosman, L., Lodder, J. C., van Ooyen, A., and Brussaard, A. B. (2005). “Role of synaptic inhibition in spatiotemporal patterning of cortical activity,” in Progress in Brain Research Progress in Brain Research. (Elsevier), 201–204. doi:10.1016/S0079-6123(04)47015-0.

Brodinová, Š., Filzmoser, P., Ortner, T., Breiteneder, C., and Rohm, M. (2019). Robust and sparse k-means clustering for high-dimensional data. Advances in Data Analysis and Classification 13, 905–932.

Carlyle, B. C., Kitchen, R. R., Kanyo, J. E., Voss, E. Z., Pletikos, M., Sousa, A. M. M., et al. (2017). A multiregional proteomic survey of the postnatal human brain. Nat Neurosci 20, 1–15. doi:10.1038/s41593-017-0011-2.

Chang, W. C. (1983). On Using Principal Components before Separating a Mixture of Two Multivariate Normal Distributions. Journal of the Royal Statistical Society: Series C (Applied Statistics) 32, 267–275. doi:10.2307/2347949.

Cody, H., Pelphrey, K., and Piven, J. (2002). Structural and functional magnetic resonance imaging of autism. Int. J. Dev. Neurosci. 20, 421–438. doi:10.1016/s0736-5748(02)00053-9.

Cooper, L. N., and Bear, M. F. (2012). The BCM theory of synapse modification at 30: interaction of theory with experiment. Nat. Rev. Neurosci. 13, 798–810. doi:10.1038/nrn3353.

Dahlhaus, M., Li, K. W., van der Schors, R. C., Saiepour, M. H., van Nierop, P., Heimel, J. A., et al. (2011). The synaptic proteome during development and plasticity of the mouse visual cortex. Mol. Cell Proteomics 10, M110.005413–M110.005413. doi:10.1074/mcp.M110.005413.

Ding, C., and Peng, H. (2005). Minimum redundancy feature selection from microarray gene expression data. J Bioinform Comput Biol 3, 185–205. doi:10.1142/s0219720005001004.

Donaldson, J., and Donaldson, M. J. (2010). Package “tsne.” Available at: https://cran.r-project.org/package=tsne.

Hastie, T., Tibshirani, R., Narasimhan, B., Chu, G., and Narasimhan, M. B. (2019). Package “impute.” CRAN Repository.

Heimel, J. A., van Versendaal, D., and Levelt, C. N. (2011). The role of GABAergic inhibition in ocular dominance plasticity. Neural Plasticity 2011, 391763–11. doi: 10.1155/2011/391763.

Hensch, T. K. (2005). Critical period plasticity in local cortical circuits. Nat. Rev. Neurosci. 6, 877–888. doi:10.1038/nrn1787.

Hensch, T. K., and Quinlan, E. M. (2018). Critical periods in amblyopia. Vis Neurosci 35, E014. doi:10.1017/S0952523817000219.

Hotelling, H., 1933 (1933). Analysis of a complex of statistical variables into principal components. Jounral of Educational Psychology 24, 417–441.

Hoyle, D. C. (2008). Automatic PCA Dimension Selection for High Dimensional Data and Small Sample Sizes. Journal of Machine Learning Research 9, 2733–2759.

Hudson, F., Josse, J., Lê, S., and Mazet, J. (2019). FactoMineR: Multivariate Exploratory Data Analysis and Data Mining. Available at: https://cran.r-project.org/web/packages/FactoMineR/.

Jolliffe, I. T., and Cadima, J. (2016). Principal component analysis: a review and recent developments. Philos Trans A Math Phys Eng Sci 374, 20150202. doi:10.1098/rsta.2015.0202.

Jones, D. G., Beston, B. R., and Murphy, K. M. (2007). Novel application of principal component analysis to understanding visual cortical development. BMC Neurosci 8, P188. doi:10.1186/1471-2202-8-S2-P188.

Lê, S., Josse, J., software, F. H. J. O. S., 2008 (2008). FactoMineR: an R package for multivariate analysis. J. Stat. Soft. 25, 1–18. Available at: https://www.jstatsoft.org/article/view/v025i01.

Luo, C., Keown, C. L., Kurihara, L., Zhou, J., He, Y., Li, J., et al. (2017). Single-cell methylomes identify neuronal subtypes and regulatory elements in mammalian cortex. Science 357, 600–604. doi:10.1126/science.aan3351.

Maaten, L. V. D., and Hinton, G. (2008). Visualizing Data using t-SNE. Journal of Machine Learning Research 9, 2579–2605.

Maffei, A., and Turrigiano, G. (2008). The age of plasticity: developmental regulation of synaptic plasticity in neocortical microcircuits. Prog. Brain Res. 169, 211–223. doi:10.1016/S0079-6123(07)00012-X.

Majdan, M., and Shatz, C. J. (2006). Effects of visual experience on activity-dependent gene regulation in cortex. Nat Neurosci 9, 650–659. doi:10.1038/nn1674.

Marini, F., and Binder, H. (2019). pcaExplorer: an R/Bioconductor package for interacting with RNA-seq principal components. BMC Bioinformatics 20, 331–8. doi:10.1186/s12859-019-2879-1.

Nowakowski, T. J., Bhaduri, A., Pollen, A. A., Alvarado, B., Mostajo-Radji, M. A., Di Lullo, E., et al. (2017). Spatiotemporal gene expression trajectories reveal developmental hierarchies of the human cortex. Science 358, 1318–1323. doi:10.1126/science.aap8809.

Revelle, W. (2019). psych: procedures for psychological, psychometric, and personality research. Northwestern University. Evanston, Illinois, R package version 1.1. Available at: https://cran.r-project.org/package=psych.

Sheng, M., Cummings, J., Roldan, L. A., Jan, Y. N., and Jan, L. Y. (1994). Changing subunit composition of heteromeric NMDA receptors during development of rat cortex. Nature 368, 144–147. doi:10.1038/368144a0.

Siu, C. R., and Murphy, K. M. (2018). The development of human visual cortex and clinical implications. Eye Brain 10, 25–36. doi:10.2147/EB.S130893.

Smith, G. B., Heynen, A. J., and Bear, M. F. (2009). Bidirectional synaptic mechanisms of ocular dominance plasticity in visual cortex. Philosophical Transactions of the Royal Society B: Biological Sciences 364, 357–367. doi:10.1098/rstb.2008.0198.

Smith, M. R., Burman, P., Sadahiro, M., Kidd, B. A., Dudley, J. T., and Morishita, H. (2016). Integrative Analysis of Disease Signatures Shows Inflammation Disrupts Juvenile Experience-Dependent Cortical Plasticity. eNeuro 3, ENEURO.0240–16.2016. doi: 10.1523/ENEURO.0240-16.2016.

Smith, M. R., Readhead, B., Dudley, J. T., and Morishita, H. (2019). Critical period plasticity-related transcriptional aberrations in schizophrenia and bipolar disorder. Schizophrenia Research 207, 12–21. doi:10.1016/j.schres.2018.10.021.

Tibshirani, R., Walther, G., and Hastie, T. (2001). Estimating the number of clusters in a data set via the gap statistic. Journal of the Royal Statistical Society: Series B (Statistical Methodology) 63, 411–423. doi:10.1111/1467-9868.00293.

Tropea, D., Kreiman, G., Lyckman, A., Mukherjee, S., Yu, H., Horng, S., et al. (2006). Gene expression changes and molecular pathways mediating activity-dependent plasticity in visual cortex. Nat Neurosci 9, 660–668. doi:10.1038/nn1689.

Warnes, G. R., Bolker, B., Bonebakker, L., Gentleman, R., Liaw, W. H. A., Lumley, T., et al. (2015). gplots: various R programming tools for plotting data. R package version 2.17. 0. Available at: https://cran.r-project.org/package=gplots.

Wei, Y., Tang, Y., Shireman, E., McNicholas, P. D., and Steinley, D. L. (2017). Extending Growth Mixture Models Using Continuous Non-Elliptical Distributions.

Wickham, H. (2009). ggplot2. New York, NY: Springer Science & Business Media.

Yashiro, K., and Philpot, B. D. (2008). Regulation of NMDA receptor subunit expression and its implications for LTD, LTP, and metaplasticity. Neuropharmacology 55, 1081–1094. doi: 10.1016/j.neuropharm.2008.07.046.

